# Simulation based benchmarking of isoform quantification in single-cell RNA-seq

**DOI:** 10.1101/248716

**Authors:** Jennifer Westoby, Marcela Sjoberg, Anne Ferguson-Smith, Martin Hemberg

## Abstract

Single-cell RNA-seq has the potential to facilitate isoform quantification as the confounding factor of a mixed population of cells is eliminated. We carried out a benchmark for five popular isoform quantification tools. Performance was generally good when run on simulated data based on SMARTer and SMART-seq2 data, but was poor for simulated Drop-seq data. Importantly, the reduction in performance for single-cell RNA-seq compared with bulk RNA-seq was small. An important biological insight comes from our analysis of real data which showed that genes that express two isoforms in bulk RNA-seq predominantly express one or neither isoform in individual cells.

## Background

RNA-seq has transformed genomics by making it easy and relatively inexpensive to obtain genome-wide, quantitative measurements of the transcriptome. To be able to take full advantage of this data; however, it is critical to have robust and accurate computational methods for quantifying transcript abundances. Isoform quantification is considered a hard problem [1–3] and the main reason why it is so challenging for mammals is because many genes have multiple isoforms which are highly similar in terms of sequence and exon structure. Because of this overlap many reads from RNA-seq experiments cannot be uniquely assigned.

Although a number of strategies have been developed to attempt to quantify isoforms [4–12], the problem of how to deal with reads which map to more than one isoform has not been fully solved [13, 14].

Although bulk RNA-seq data of cells indicate the expression of multiple isoforms, scRNA-seq of the same cells show evidence for a single or a small number of isoforms per gene [15–18]. This is potentially beneficial, as it suggests that by performing isoform quantification using single-cell RNA-seq (scRNA-seq) data rather than bulk RNA-seq data, the problem could be simplified due to a reduced number of multi-mapping reads. As well as allowing basic biological questions on cellular heterogeneity to be investigated, isoform quantification using scRNA-seq could have applications in fields such as cancer, developmental biology, alternative splicing and neurobiology.

Despite the intrinsic benefits of single-cell resolution, to date most scRNA-seq research has focused on gene level quantification and analysis [19]. This partly reflects the novelty of single-cell RNA sequencing technologies and best practices for performing isoform quantification using single cell data have yet to be established. Numerous tools to perform isoform quantification are available [4–12], but most of these tools were designed for bulk RNA-seq analysis and it is not clear whether it would be appropriate to apply these tools to single-cell analysis. For other tasks, e.g. normalization, It has been shown that applying methods designed for bulk RNA-seq to scRNA-seq can give misleading results [20]. It has not yet been established whether isoform quantification methods designed for bulk RNA-seq are accurate for single-cell data.

In addition to software concerns, there are experimental and technical issues which could impact on isoform quantification in single cells. A wide range of library preparation protocols have been developed for scRNA-seq [21–29], some of which are likely to be more appropriate for isoform quantification than others. For example, one way in which library preparation protocols could differ in their suitability for isoform quantification is in their degree of gene length bias, which has been shown to be greater for full length transcript protocols compared with UMI based protocols [30]. An understanding of which library preparation protocols generate data suitable for isoform quantification and which library preparation protocols do not would allow researchers to better design experiments to suit their needs. Due to the low amount of starting material, scRNA-seq data has greater variability and a greater number of transcripts for which zero reads are detected, i.e. dropouts, relative to bulk RNA-seq data [31]. It is known that whilst some of this variation is biological in origin, a substantial proportion is technical [17, 32, 33]. Dropouts, defined as events in which reads mapping to a gene or isoform are detected in some cells but not others, are highly prevalent in scRNA-seq [34]. The impact of dropouts and other technical noise on isoform quantification tools is not known and different strategies than the ones that were used for bulk RNA-seq may be required.

To assess isoform quantification for scRNA-seq, we present a simulation based benchmarking study using data generated from three different scRNA-seq projects. Whilst benchmarking studies have been performed previously for bulk RNA-seq [13, 14], to the best of our knowledge this is the first benchmark of isoform quantification performed for scRNA-seq. We evaluated the overall accuracy of different isoform quantification methods when applied to scRNASeq, and we also specifically studied the impact of library preparation protocol and dropouts. We tested five popular isoform quantification tools on simulated scRNA-seq data based on three publicly available scRNA-seq datasets produced using different library preparation protocols and cell types. With the exception of eXpress, performance was generally good for simulated data based on SMARTer and SMART-seq2 [27] data, but poor for simulated data based on Drop-seq [23] data. Compared to bulk RNA-seq, isoform quantification was only slightly worse for SMARTer and SMART-seq2 data, suggesting that it is appropriate to use these methods for full-transcript single-cell data.

## Results

### The performance of isoform quantification tools was generally good and consistent across two different simulation methods

The first dataset considered in this benchmark consisted of 96 mouse quiescent B lymphocytes collected as part of the BLUEPRINT epigenome project [35] (GEO accession code GSE94676). The SMARTer library preparation protocol was used to collect this dataset, which has been shown to have a degree of 3’ coverage bias [36]. On average, just over 2.7 million reads had been sequenced per B lymphocyte.

To perform the benchmark, simulated data was generated from the selected cells using two simulation methods. The first simulation method used was RSEM [4] (see Methods for details). RSEM is an isoform quantification tool which uses a generative model and expectation maximisation to estimate isoform expression. In addition, RSEM is capable of simulating reads using its generative model and input values for the latent variables in the model, which can be estimated during isoform quantification. An important reason for selecting RSEM to perform the simulations is that during the simulation process, RSEM records where each simulated read originated in the transcriptome. Consequently, it is known how highly expressed each isoform is in the simulated data. This will be referred to as the ‘ground truth’. Knowing the ground truth allows us to benchmark expression estimates from isoform quantification tools using the simulated data.

The second simulation method relied on two tools, Splatter [37] and Polyester [38]. Splatter is a simulation tool which takes an expression matrix of counts from an scRNA-seq experiment as input and gives a simulated expression matrix of counts as output. Splatter was used to simulate counts data based on an expression matrix of counts from the BLUEPRINT B lymphocytes generated by Kallisto [6]. Since the exact origin in the transcriptome is not known from Splatter, the simulated expression matrix of counts was then given as input to Polyester, which generated simulated reads. The Splatter counts matrix was converted to a matrix of TPM values, which were used as the ‘ground truth’.

The RSEM and Splatter and Polyester simulated reads data was then given as input to RSEM, eXpress [5], Kallisto, Salmon [7] and Sailfish [8]. The isoform quantification tools provide two useful pieces of information for each isoform – whether it is expressed and its expression level. To quantify the ability of each method to detect the presence of an isoform, the precision and recall were calculated. In this context, the precision is the fraction of isoforms predicted to be expressed by each tool which are expressed in the ground truth. The recall is the fraction of isoforms expressed in the ground truth which are predicted to be expressed using the tool. For a single overall quality score we used the F1 score, which is defined as the harmonic mean of precision and recall.

Salmon can be run in three modes – an alignment based mode, in which aligned reads are taken as input, or one of two alignment free modes (a quasi mode or an SMEM mode). The performance of all three modes was evaluated in this benchmark. For most isoform quantification tools, the mean F1 score was remarkably similar and in the range of 0.758 - 0.885. The exception was eXpress, which had a slightly higher recall but a much lower precision than other tools, and consequently had the lowest mean F1 score (between 0.452 and 0.486 depending on simulation method) (Figure 1A). The mean F1 scores, precisions and recalls calculated for each of these tools were similar regardless of whether RSEM or Splatter and Polyester were used to generate the simulated data. The statistics were not dramatically altered when Polyester simulated reads using a 3’ coverage bias model compared to when Polyester simulated reads uniformly across transcript length. However, as the Polyester 3’ coverage bias model is not based on single cell RNA-seq data, care needs to be taken when interpreting this result.

**Figure 1.**
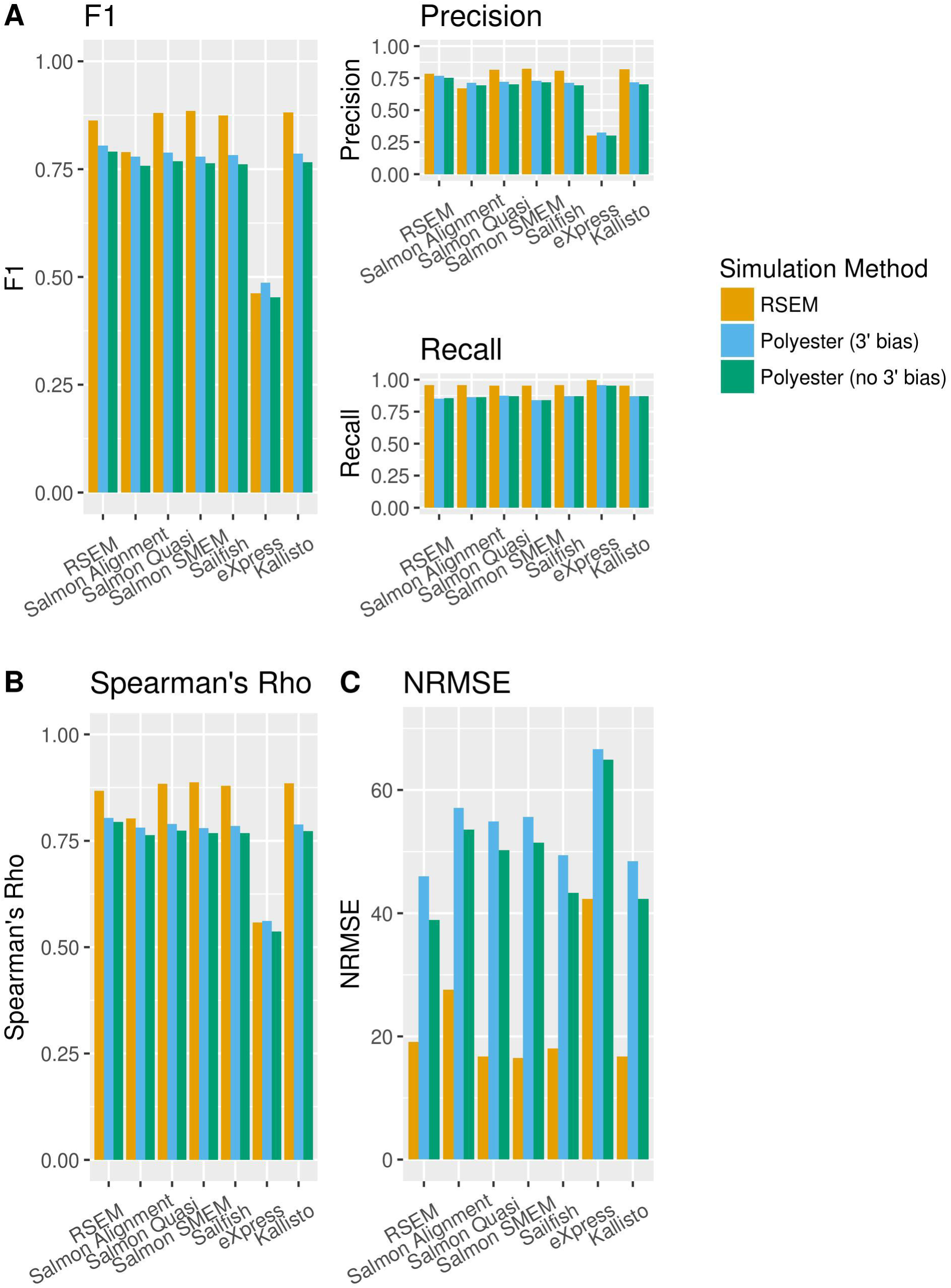
Performance statistics for each isoform quantification tool for the BLUEPRINT simulations. The yellow bars represent RSEM simulations, the blue bars represent Splatter and Polyester simulations with 3’ coverage bias and the green bars represent Splatter and Polyester simulations with no coverage bias. (**a**) F1 score, precision and recall of isoform detection. The F1 score is the harmonic mean of the precision and recall. The precision is the proportion of the isoforms predicted to be expressed by an isoform quantification tool which are expressed. The recall is the proportion of expressed isoforms which are predicted to be expressed by the isoform quantification tool. (**b**) Spearman’s rho. (**c**) Normalised Root Mean Square Error (NRMSE).

In addition to determining whether an isoform is expressed, it is often of interest to estimate isoform abundance. To evaluate how well isoform quantification tools perform this task, two measures were considered – Spearman’s rho and the normalized root mean square error (NRMSE) (Figure 1B,C). Spearman’s rho gives a measure of how monotonic the relationship between the ground truth expression and each tool’s expression estimates is, whilst the NRMSE gives a measure of the extent to which the relationship deviates from a one to one linear relationship (See Methods for a details on how the NRMSE was calculated).

Consistent with the results for isoform detection, the mean Spearman’s rho was similar between isoform quantification tools and simulation methods and in the range 0.763 - 0.888. The exception was eXpress, which had a much lower mean Spearman’s rho than the other tools with values from 0.538 to 0.562. eXpress also performed poorly relative to the other isoform quantification tools when considering the NRMSE. Interestingly, although the overall pattern of NRMSE results was similar for both simulation methods, the NRMSE was consistently far higher for the Splatter and Polyester simulations compared to the RSEM simulations. One possible explanation is that the difference in the NRMSE is due to a small number of outliers. However, this did not appear to be the case (see Figure S6). Another explanation for the difference in the NRMSE could be that the differences are largely driven by differences in the ground truth expression distributions of the RSEM simulations compared to the Splatter and Polyester simulations. Since the NRMSE is proportional to the sum of squared differences between the ground truth and the isoform quantification tool’s expression estimates, it is plausible that it will be relatively rare for an unexpressed isoform to have an estimated expression other than zero, but relatively common for an expressed isoform to have an estimated expression that differs from the ground truth expression. We found that the distribution of ground truth expression values differs for each simulation method (See Figure S7). Therefore, differences in the ground truth expression distributions seem to be the most likely explanation for the systematic difference in the NRMSE between simulation methods.

The difference in the NRMSE between simulation methods was not the only aspect in which the simulation methods differed. A comparison of the simulated data with the real data was carried out using both a comparison tool included in Splatter and using CountsimQC [39], a package which facilitates comparison of simulated datasets. The RSEM simulated data more closely resembled the real data than the Splatter and Polyester simulated data by a number of metrics, including the sample-sample correlations, the mean-variance relationship and the relationship between magnitude of expression and fraction of zeros (Figure S8). An additional way in which the Splatter and Polyester simulations differ from the real data is in terms of their transcriptional profile. Isoform names are lost in the Splatter simulation process and prior to the Polyester read simulation, simulated counts are reassigned to isoforms known to be expressed in the real data (see Methods). Consequently the correlation between ground truth isoform expression in the Splatter and Polyester simulated data and isoform expression estimates generated by running Kallisto on the real BLUEPRINT B lymphocyte data is very low (see Figure S9). In contrast, the correlation between ground truth expression in the RSEM simulations and Kallisto expression estimates in the real data was much higher. Based on these findings, we concluded that the RSEM simulations resembled the real data more closely than the Splatter and Polyester simulations. Consequently, for the rest of this paper all data was simulated using RSEM.

Despite the differences between the RSEM and Splatter and Polyester simulations, the results of the benchmark were remarkably consistent. This suggests that the findings in this benchmark are robust to some differences between datasets, including dramatic changes in the transcriptional profile.

### Isoform quantification tools generally perform well on SMART-seq2 data with high sequencing coverage

There is currently a trade-off in scRNA-seq protocols between the number of reads sequenced per cell and the number of cells sequenced [31]. An interesting question with regards to isoform quantification and detection is whether sequencing more reads per cell or sequencing more cells is more effective. Whilst sequencing at low read coverage is likely to reduce the accuracy of isoform quantification and detection in individual cells, it is possible that this loss of information could be compensated to some extent by sequencing a higher number of cells. To investigate the impact of the trade-off between the number of reads sequenced per cell and the number of cells sequenced on the performance of isoform quantification tools, two datasets with a very different number of reads per cell and a very different total number of cells were considered.

The first dataset considered was a mouse embryonic stem cells (mESCs) dataset published by Kolodziejczyk et al. [40]. On average, over 7 million reads were sequenced per cell in this dataset. From this dataset, 271 cells mESCs grown in standard 2i media + LIF which passed quality control were used for the benchmark (see Methods). This dataset should therefore give a good indication of the performance of isoform quantification tools when there are a high number of reads per cell but a relatively low number of cells. In addition, this dataset has uniform coverage of transcripts, as it was sequenced using the SMART-seq2 protocol [27].

To perform the benchmark, simulated data was generated as described previously from the selected cells from Kolodziejczyk et al. (see Methods for details). The simulated reads data were then given as input to RSEM, eXpress, Kallisto, Salmon and Sailfish. The highest F1 score was achieved by Salmon run in SMEM mode (0.889), with RSEM, Salmon run in quasi mode, Sailfish and Kallisto also achieving mean F1 scores greater than 0.85 (Figure 2A). Again, eXpress performed most poorly by a substantial margin, with a mean F1 score of 0.548, and again, eXpress had a higher mean recall (0.997) but a much lower mean precision (0.378) than other tools. It seems likely that eXpress’s low precision is due to it being too liberal when calling isoforms as expressed. The average number of isoforms called as expressed per cell was twice as high for eXpress, which called an average of 41,339 isoforms as expressed per cell, as for any other tool. The other isoform quantification tools had high mean recalls between 0.956 and 0.960. In contrast, the highest mean precision was just 0.831 by Salmon run in SMEM mode, which means that nearly one in six isoforms predicted to be expressed by the best performing tool were not actually expressed. The high recall values achieved by all the tools considered here indicates that the vast majority of isoforms expressed in the simulated data are detected, with the lower precision values being a greater cause for concern. Knowing that an isoform is not expressed can be as important as knowing that an isoform is expressed, especially if that isoform is being used as a marker, for example in clustering analysis. A strategy for improving the detection ability as quantified by the F1 score of isoform quantification tools for scRNA-seq could be to make future tools more conservative when calling isoforms as expressed.

**Figure 2.**
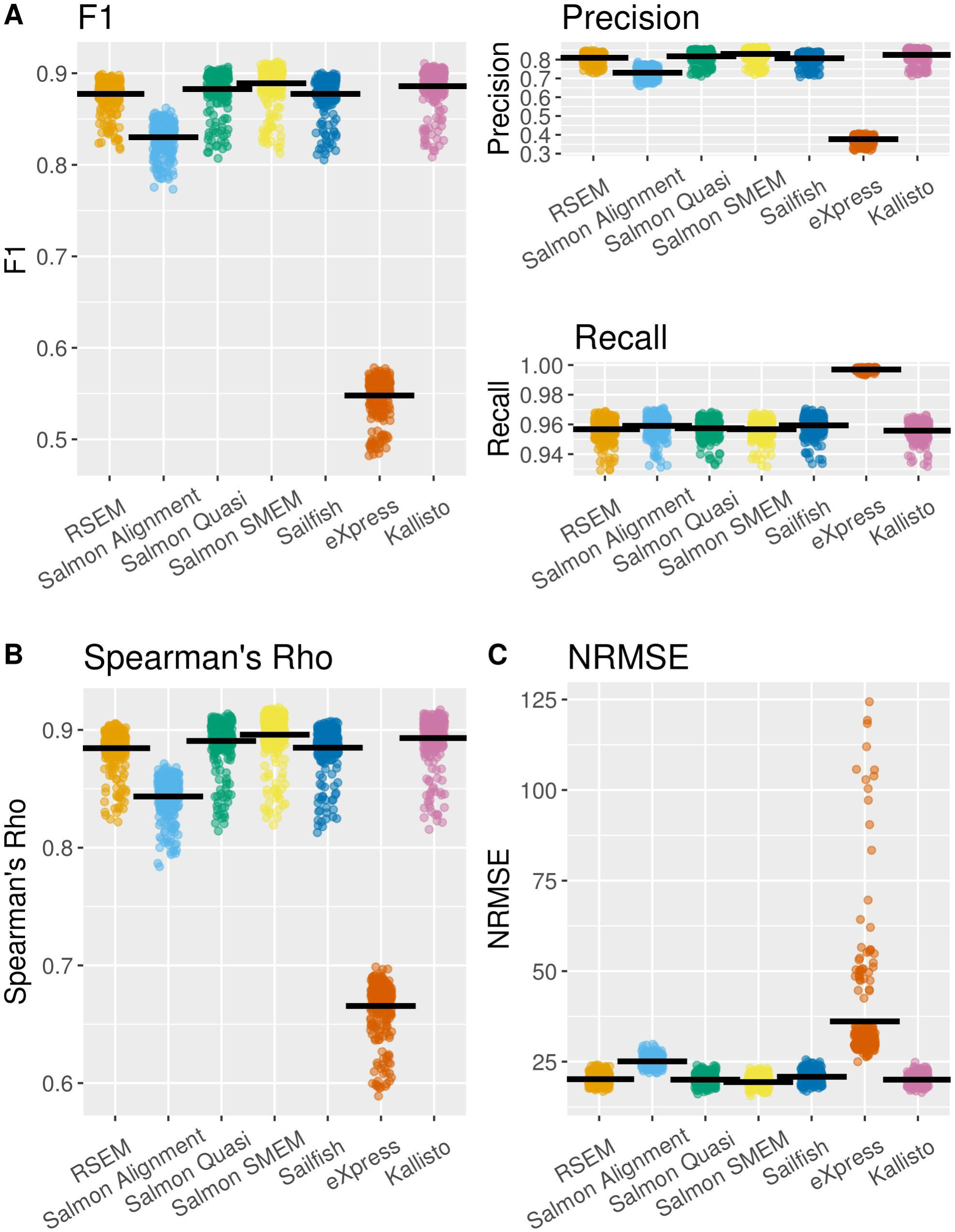
Performance statistics for each isoform quantification tool for the BLUEPRINT simulations. The yellow bars represent RSEM simulations, the blue bars represent Splatter and Polyester simulations with 3’ coverage bias and the green bars represent Splatter and Polyester simulations with no coverage bias. (**a**) F1 score, precision and recall of isoform detection. The F1 score is the harmonic mean of the precision and recall. The precision is the proportion of the isoforms predicted to be expressed by an isoform quantification tool which are expressed. The recall is the proportion of expressed isoforms which are predicted to be expressed by the isoform quantification tool. (**b**) Spearman’s rho. (**c**) Normalised Root Mean Square Error (NRMSE).

The highest mean value of Spearman’s rho was obtained by Salmon run in SMEM mode (0.896), with Salmon run in quasi mode, Kallisto, RSEM and Sailfish obtaining similar values.

The lowest mean value of the NRMSE was also obtained by Salmon run in SMEM mode (19.5), with Salmon run in quasi mode, Kallisto, RSEM and Sailfish obtaining similar values. Again, of the tools considered, eXpress performed most poorly by a substantial margin.

### The performance of isoform quantification tools was generally poor using the Drop-seq library preparation method

Droplet based library preparation methods for scRNA-seq enable tens or hundreds of thousands of cells to be sequenced in a single experiment, but at a relatively low coverage per cell [23, 24, 26, 41]. To determine whether a high number of cells can compensate for low sequencing depth, a Drop-seq dataset of retinal bipolar cells published by Shekhar et al. [42] was considered. Approximately 45,000 cells were sequenced at a median read depth of 8,200 mapped reads per cell in this dataset. From the dataset, 1,000 cells were randomly selected and given as input to RSEM to generate simulated data (see Methods for details). The simulated reads were then given as input to RSEM, eXpress, Kallisto, Salmon and Sailfish as before, and the performance of these tools for the Drop-seq simulated data was evaluated.

With the exception of RSEM, the mean F1 score is far lower for the Shekhar et al. Drop-seq simulated data as compared with the Kolodziejczyk et al. or the BLUEPRINT simulated data (Figure 3A). For most tools, this is a consequence of a drop in both the precision and the recall. The mean precision is less than 0.5 for most isoform quantification tools, including those which performed well on the Kolodziejczyk et al. and BLUEPRINT simulated data. Salmon run on SMEM mode, which achieved the highest mean precision on the Kolodziejczyk et al. simulated data, performed particularly poorly, achieving a mean precision of just 0.399. Only RSEM and Kallisto achieved mean precisions greater than 0.5 (0.743 and 0.614 respectively).

**Figure 3.**
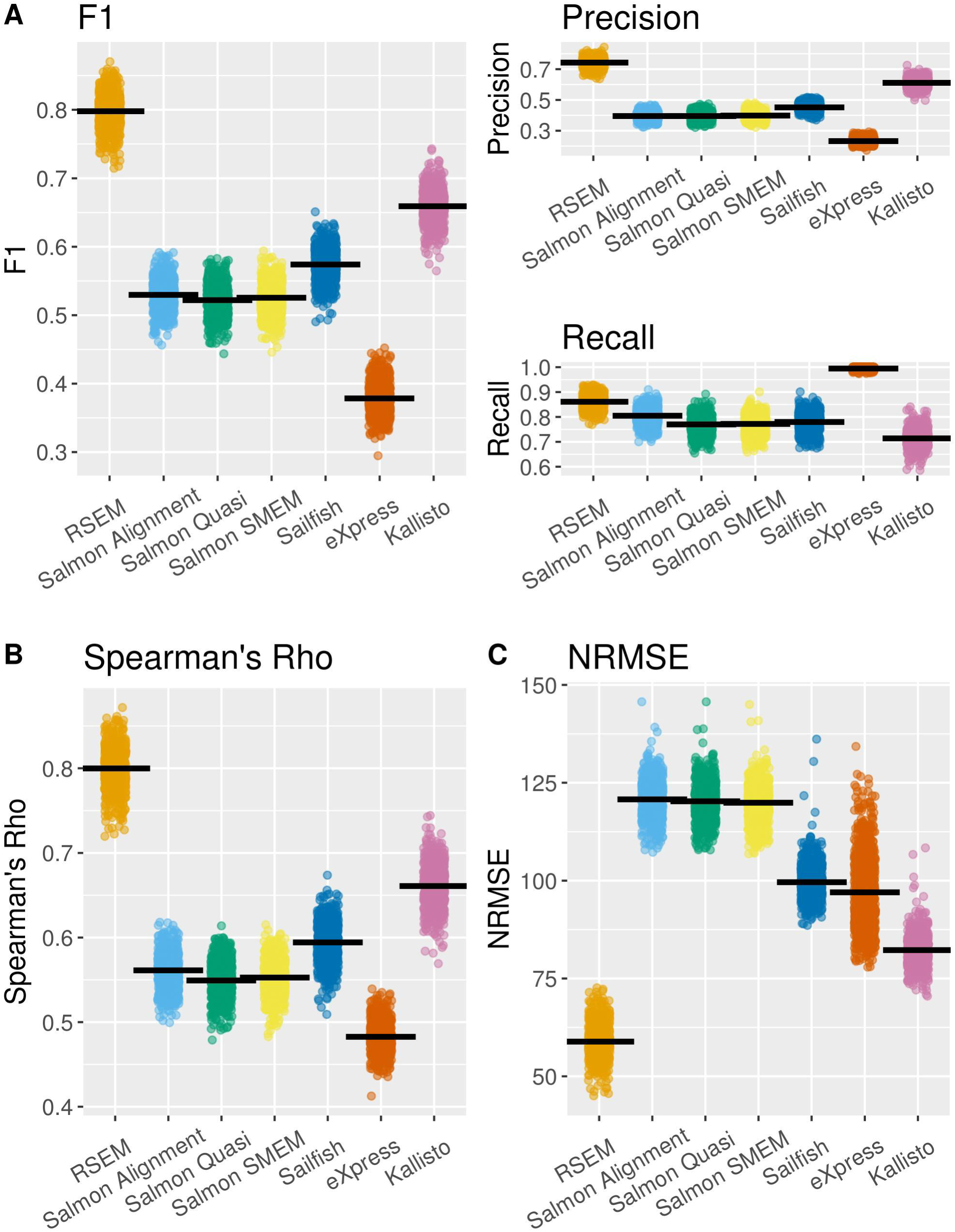
Performance statistics for each isoform quantification tool for the Shekhar et al. Drop-seq simulations. (**a**) F1 score, precision and recall of isoform detection. The F1 score is the harmonic mean of the precision and recall. The precision is the proportion of the isoforms predicted to be expressed by an isoform quantification tool which are expressed. The recall is the proportion of expressed isoforms which are predicted to be expressed by the isoform quantification tool. (**b**) Spearman’s rho. (**c**) Normalised Root Mean Square Error (NRMSE).

This result has important implications for the notion that sequencing a high number of cells could capture the overall transcriptional profile of a population of cells despite a low number of reads per cell. If sequencing a high number of cells could compensate for a low coverage per cell, a very high precision of isoform detection would be required. Without a high precision, attempting to combine data from multiple cells to recapture the population transcriptional profile will result in calling a high number of unexpressed isoforms as expressed, whereas if low read numbers only reduced recall it could be compensated for by combining data across cells. In addition to a low precision and recall, the isoform quantification tools perform relatively poorly on the Shekhar et al. Drop-seq simulated data when Spearman’s rho and NRMSE are used as performance metrics (Figure 3B,C). Again, Kallisto and RSEM perform relatively well by these metrics compared to the other tools. The overall picture painted by these results is that a low number of reads per cell reduces the performance of isoform quantification tools, and this cannot be compensated for by sequencing more cells.

RSEM appears to perform better than the other isoform quantification tools when run on the Shekhar et al. Drop-seq simulated data, however this result needs to be interpreted with caution. Since RSEM was used to perform the Shekhar et al. Drop-seq simulations, and as it essentially uses the same model to perform isoform quantification and simulations, it is plausible that RSEM’s performance is close to optimal when run on its own simulated data. This was also the case for the Kolodziejczyk et al. and some of the BLUEPRINT simulations, but for these simulations the performance of Sailfish, Salmon and Kallisto was not dissimilar to the performance of RSEM. One hypothesis generated from these observations is that on high quality single cell datasets, most isoform quantification tools perform well, meaning that if RSEM is performing optimally, it provides a relatively small advantage. However, on a dataset with a low number of reads per cell, short reads and 3’ coverage bias, most tools perform poorly. If RSEM is performing optimally, this may result in a much greater impact on relative performance.

### The decrease in the performance of isoform quantification using scRNA-seq compared with bulk RNA-seq is generally small

We find that the performance of existing isoform quantification tools is generally good when run on simulated data based on SMART-seq2 and SMARTer scRNA-seq data. We next consider the performance of isoform quantification tools when scRNA-seq data is used compared with bulk RNA-seq data. Although previous benchmarks of isoform quantification have been performed using bulk RNA-seq data [13, 14], a direct comparison with our benchmark is challenging due to differences in the experimental approaches taken. Consequently, it is not possible to say whether any perceived change in the performance of a given tool in our benchmark compared with a bulk RNA-seq benchmark is due to differences in how the benchmark was performed, differences in which statistics were collected, or due to a genuine difference in performance on bulk and single-cell data.

To gain further insights regarding the performance of the tools, we made use of the bulk RNA-seq data generated for the BLUEPRINT B lymphocytes and Kolodziejczyk et al. standard 2i media + LIF mESCs. We used RSEM to simulate the bulk RNA-seq data and collected the same performance statistics for our bulk RNA-seq benchmark as in our scRNA-seq benchmark. As the data used in our bulk and scRNA-seq benchmark came from the same source, the same method was used to generate the simulated bulk and scRNA-seq data, and the same performance statistics were collected in both benchmarks, we were able to perform a meaningful comparison of isoform quantification tool performance on bulk and scRNA-seq data.

We find that all isoform quantification tools performed well on the simulated bulk-data, but since most methods also performed well on single-cell data the improvement was generally small (Figure 4 and S16). In particular, there is very little difference in the recall for bulk and scRNA-seq, for which performance seems to be close to optimal. Interestingly, eXpress performs far better on bulk RNA-seq compared with scRNA-seq. Since eXpress appears to be overly liberal in calling isoforms as expressed, one explanation for the better performance of eXpress on bulk RNA-seq is that more isoforms have non-zero expression in bulk (Supplementary Figure S17). Consequently, there are fewer unexpressed isoforms for eXpress to incorrectly call as expressed.

**Figure 4.**
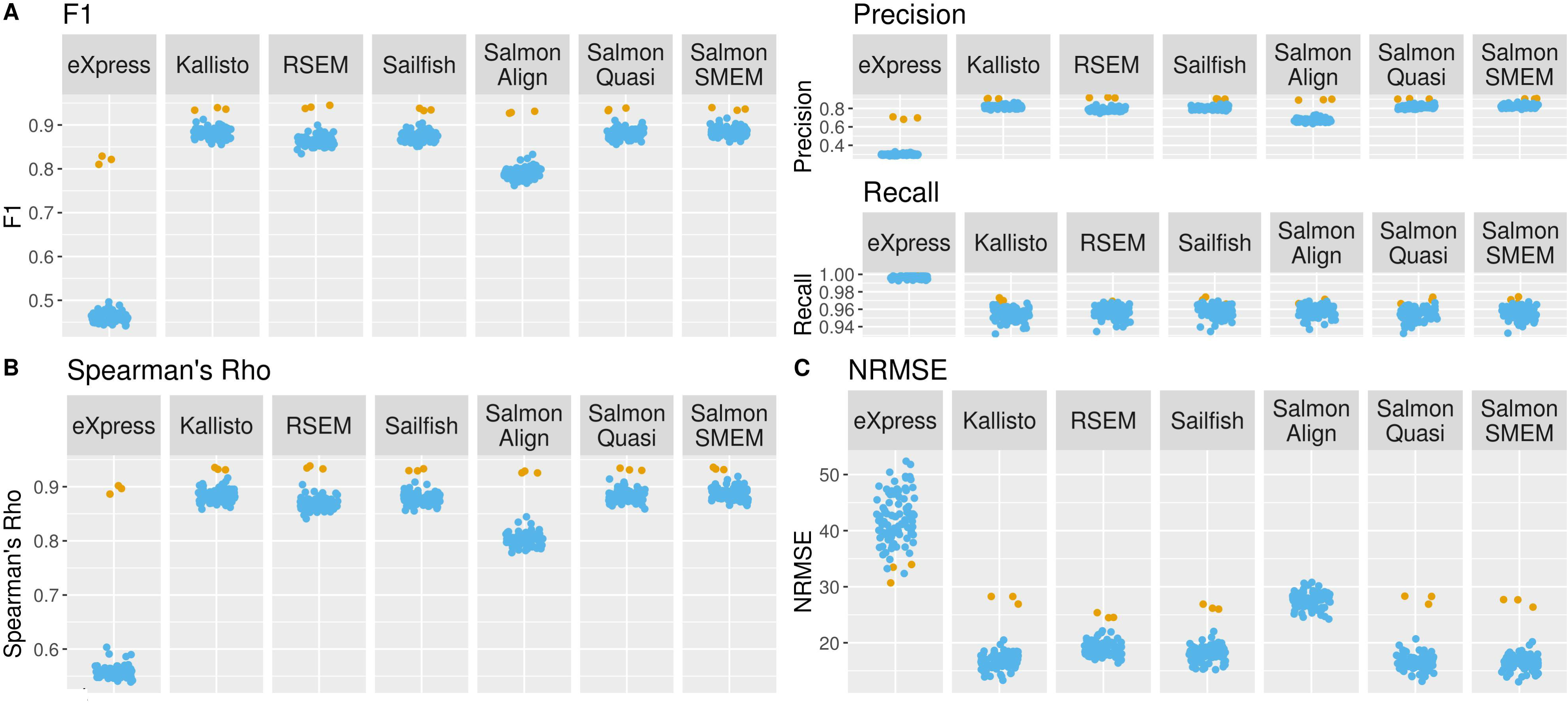
Comparison of the performance of isoform quantification tools on BLUEPRINT B lymphocyte bulk and single cell RNA-seq data. Each point represents one cell from the scRNA-seq dataset or one bulk RNA-seq experiment. Yellow points represent bulk RNA-seq experiments, blue points represent one cell from the scRNA-seq experiment. (**a**) F1 score, precision and recall of isoform detection. The F1 score is the harmonic mean of the precision and recall. The precision is the proportion of the isoforms predicted to be expressed by an isoform quantification tool which are expressed. The recall is the proportion of expressed isoforms which are predicted to be expressed by the isoform quantification tool. (**b**) Spearman’s rho. (**c**) Normalised Root Mean Square Error (NRMSE).

Salmon run on alignment mode performs surprisingly poorly on the simulated Kolodziejczyk et al. bulk RNA-seq data (Figure S16), despite performing far better on the BLUEPRINT bulk RNA-seq data and on both single cell datasets. As there was only one standard 2i media + LIF mESC bulk RNA-seq dataset, it is not clear whether this is an outlier or whether this result would have been robust had additional standard 2i media + LIF mESC bulk RNA-seq replicates been collected. However, the result suggests that the performance of Salmon run on alignment mode can be variable when run on bulk RNA-seq.

### Removing drop-outs can improve the performance of isoform quantification tools

Whilst it is of interest to determine which isoform quantification tools perform best overall when run on scRNA-seq data, it is important to recognise that such an analysis may hide a lot of detail. For example, scRNA-seq data commonly contains a high number of dropouts [43], and one question of interest is whether the performance of isoform quantification tools differs between isoforms with a high number of dropouts and isoforms with few or no dropouts.

To address the impact of dropouts on performance, Spearman’s rho and the NRMSE were calculated when isoforms with zero expression in more than a specific fraction of cells were removed from the analysis. Interestingly, applying increasingly stringent thresholds to remove isoforms with a high number of dropouts led to an increase in the value of Spearman’s rho in both the Kolodziejczyk et al. and BLUEPRINT simulations (Figure 5a). For isoforms which had dropouts in less than 20% of cells, the value of Spearman’s rho became very high for Sailfish, Salmon, Kallisto and RSEM (in the range of 0.992-0.997 for the BLUEPRINT simulations, and 0.977-0.989 for the Kolodziejczyk et al. simulations). This indicates that for isoforms with very few dropouts, isoform quantification tools are extremely good at ordering their relative expression correctly. Removing isoforms with a high number of dropouts had a more variable effect on the NRMSE. Due to the inverse relationship between magnitude of expression and number of dropouts [34] in both the real and simulated data (Figure 5b), one explanation for the increase in Spearman’s rho is that lowly expressed isoforms are more likely to have a high number of dropouts and are also more likely to be mis-ordered with respect to the ground truth. However, because they are lowly expressed, removing them has a relatively small effect on the NRMSE.

**Figure 5.**
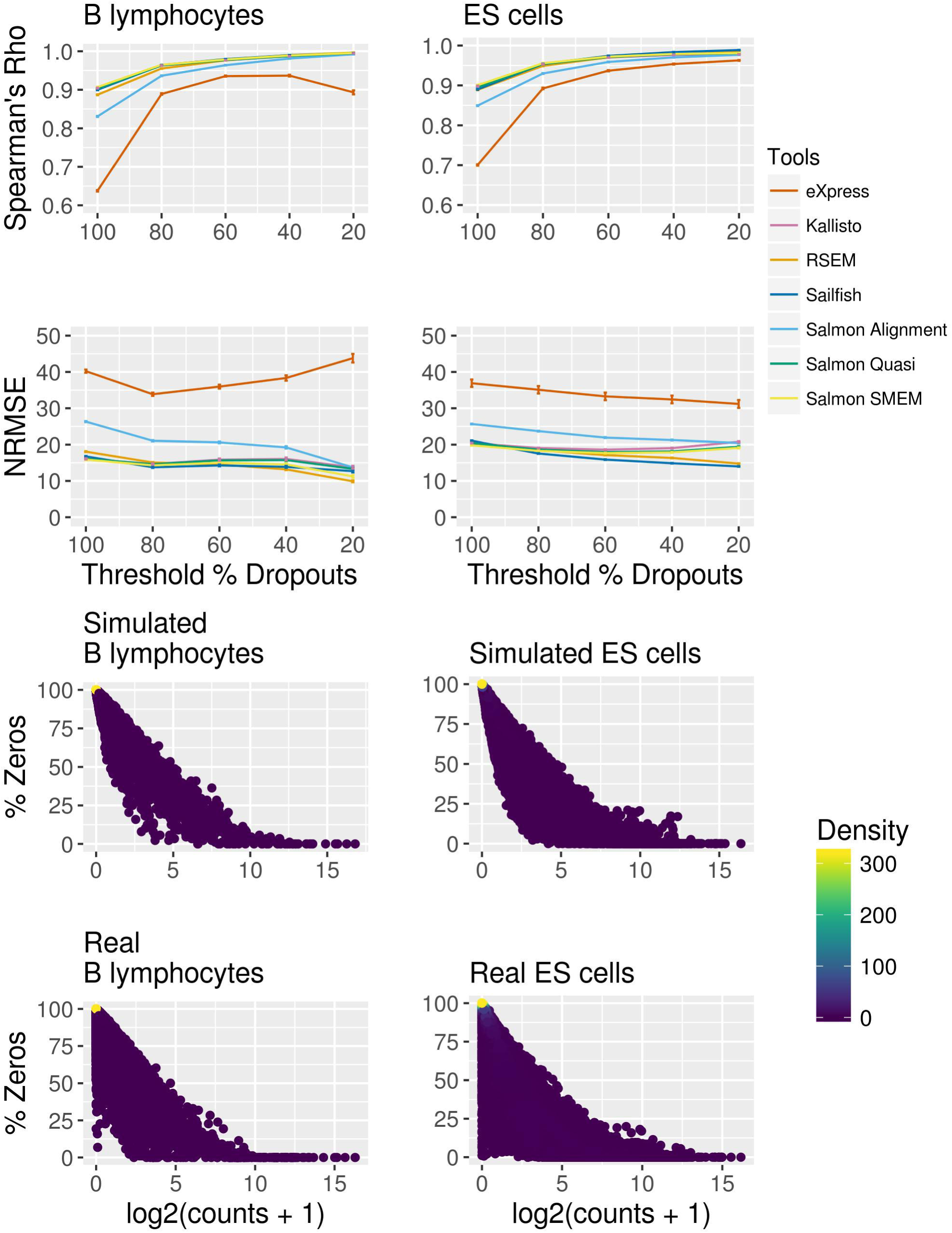
Effect of dropouts on isoform quantification. (**a**) Impact of removing isoforms with more than a threshold number of dropouts on Spearman’s rho and the NRMSE for the BLUEPRINT B lymphocytes (left) and the Kolodziejczyk et al. ES cells (right). The x axis gives the threshold percentage of zeros above which an isoform is removed from the analysis. For example, a threshold percentage of 80% would result in isoforms with zero expression in 80% or more of cells being removed from the analysis. Each coloured line is a linear fit for visual guidance and it represents a different isoform quantification tool. (**b**) Relationship between how highly expressed an isoform is and the percentage of cells in which it has zero expression. The relationship is considered in RSEM simulated BLUEPRINT B lymphocytes (top left), the real BLUEPRINT B lymphocytes (bottom left), the simulated Kolodziejczyk et al. ES cells (top right) and the real Kolodziejczyk et al. ES cells (bottom right). Each point represents an isoform and the points are coloured according to density.

### For genes which express two isoforms in bulk RNA-seq, usually only one isoform is detected per cell in scRNA-seq

To determine whether individual cells express all or only some of the isoforms seen in a population of cells, we consider genes which have two isoforms, both of which are expressed in the BLUEPRINT B lymphocyte or in the Kolodziejczyk et al. ES cell bulk RNA-seq data. We then determine how many isoforms are expressed from these genes in the corresponding scRNA-seq data. Kallisto was used to perform isoform quantification for the bulk and single cell data as it performed well in both our bulk and single cell benchmarks.

For genes which express two isoforms in bulk RNA-seq data, we first consider if zero, one or two isoforms are detected in single cells. For most genes which express two isoforms in the bulk RNA-seq, neither isoform is expressed in most cells in the scRNA-seq (Figure 6A). It is more common in cells which do express the gene to express one rather than two isoforms. To investigate this further, we consider the percentage of cells which express both isoforms expressed in the bulk RNA-seq. We find that for the majority of genes, no or very few cells express both isoforms seen in the bulk RNA-seq, however for a minority of genes in both the BLUEPRINT B lymphocytes and Kolodziejczyk et al. ES cells, a high percentage of cells express both isoforms (Figure 6B). Interestingly, more genes express both isoforms in the Kolodziejczyk et al. ES cells compared to the BLUEPRINT B lymphocytes. This may partly reflect the higher number of cells and the higher number of reads per cell in the Kolodziejczyk et al. ES cells, possibly enabling better detection of lowly and/or infrequently expressed isoforms.

**Figure 6.**
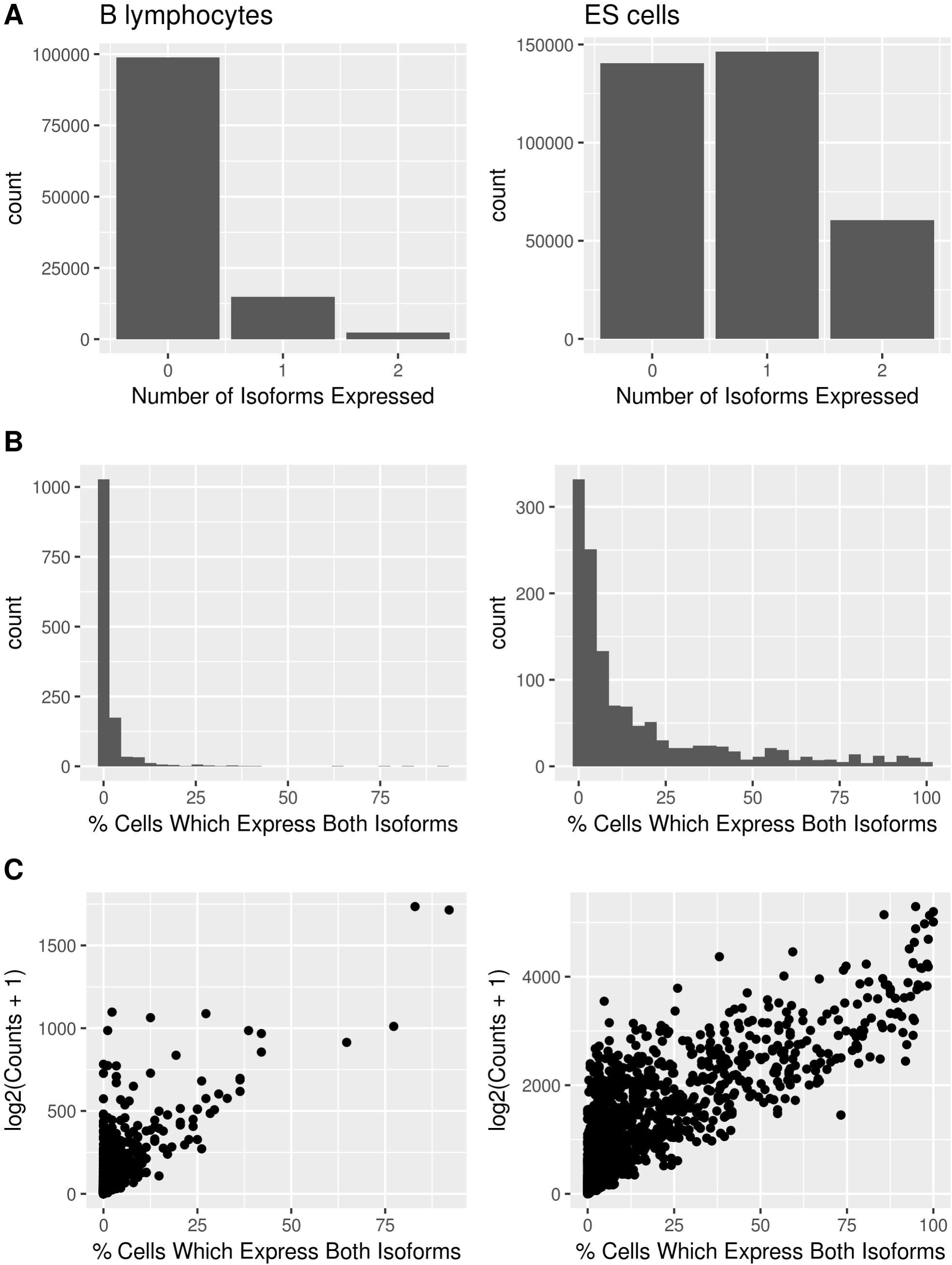
Investigation into how many isoforms are expressed per cell in the scRNA-seq data for genes which express exactly two isoforms. The BLUEPRINT B lymphocyte (left) and the Kolodziejczyk et al. ES cell (right) bulk RNA-seq data are shown. The B lymphocyte graphs shown here are from the first biological replicate of the BLUEPRINT male B lymphocyte bulk RNA-seq, equivalent graphs for the second and third BLUEPRINT male B lymphocyte biological replicates can be found in Figure S18 (**a**) Number of genes which express two isoforms in the bulk RNA-seq data expressing zero, one or two isoforms in each cell in the scRNA-seq data. (**b**) Histogram of the percentage of cells which express both the isoforms detected in the bulk RNA-seq data. (**c**) Relationship between the percentage of cells which express both the isoforms detected in the bulk RNA-seq. Spearman’s rho is 0.623 for the BLUEPRINT B lymphocytes and 0.795.for the Kolodziejczyk et al. ES cells.

In addition, the globally elevated transcription rates in ES cells relative to other cell types might lead us to expect that expression of multiple isoforms from a single gene would be more common in ES cells [44]. Finally, we ask whether more highly expressed genes are more likely to express multiple isoforms. We find a positive correlation between gene expression and the percentage of cells which express both isoforms (Figure 6C), indicating that more highly expressed genes are more likely to express both isoforms in individual cells.

## Discussion

To date, scRNA-seq studies have mainly focussed on gene level quantification [19]. This has partly been due to uncertainty over how best to perform isoform quantification in scRNA-seq. In addition, there has been uncertainty over whether the results obtained would be meaningful due to the low read coverage compared with bulk RNA-seq. Our analyses have demonstrated that Kallisto, Salmon, Sailfish and RSEM can accurately detect and quantify isoforms in scRNA-seq to nearly the same accuracy as bulk RNA-seq data, provided the datasets have a reasonably high number of reads per cell. For simulated data based on Drop-seq, the performance of isoform quantification tools was too poor to make the results of performing isoform quantification worthwhile. Due to the low precision of isoform detection, this problem cannot be overcome by incorporating information from a large number of cells. It is possible that increasing the number of reads sequenced per cell for Drop-seq protocols would improve the performance of isoform quantification tools, although the short reads and 3’ coverage bias are likely to ensure that accurate quantification remains challenging.

In addition to benchmarking isoform quantification for scRNA-seq, we perform an equivalent benchmark for bulk RNA-seq. We find that the performance of most isoform quantification tools is slightly worse for scRNA-seq compared with bulk, but that the difference is small. The cost in performance using scRNA-seq compared with bulk RNA-seq for isoform quantification is therefore low. However, it should be noted that this benchmark has evaluated the ability of isoform quantification tools to correctly assign the reads present in an scRNA-seq experiment to the isoforms they originated from. As a major technical issue with scRNA-seq is failure to capture reads from a high proportion of expressed transcripts [20], it is likely that in practice, many expressed isoforms will be missed by isoform quantification tools when run on scRNA-seq data due to a lack of evidence in the captured reads that the isoform was expressed. However, the extremely high recall of all the isoform quantification tools considered in this benchmark means that the overwhelming majority of isoforms from which reads are captured will be called as expressed. More problematic is the relatively low precision of isoform detection, as a consequence of which around one in six isoforms called as expressed in deeply sequenced scRNA-seq datasets will be false positives, even for the best performing tools.

Whilst our analysis has demonstrated that existing tools can accurately detect and quantify isoforms for scRNA-seq, no tool performed perfectly. The tools benchmarked here were designed for use with bulk RNA-seq, and it is plausible that future tools designed to perform isoform quantification specifically for scRNA-seq could perform better. We found that the tools benchmarked in this study tended to have a higher recall than precision of isoform detection. Therefore, it is likely the performance of isoform quantification tools designed for scRNA-seq data could be improved by making the tools more conservative in calling isoforms as expressed relative to tools designed for use on bulk data. In addition, we found that Spearman’s rho increased when lowly expressed isoforms with a high number of dropouts were removed from the analysis. Thus, it is likely that attempts to incorporate the effects single cell specific technical noise such as dropouts would improve the performance of isoform quantification tools on scRNA-seq. An open question for isoform quantification in scRNA-seq is whether incorporating information from Unique Molecular Identifiers (UMIs) into isoform expression estimates could improve accuracy of quantification. Whilst UMI information could reduce the effects of PCR amplification noise [45], UMI based protocols tend to exhibit significant coverage bias, potentially making isoform quantification challenging [46]. If UMI based protocols could be combined with long read sequencing technologies, this problem could potentially be overcome.

A number of previous studies have attempted to determine whether genes from which multiple isoforms are expressed in a population of cells are expressing all of these isoforms in individual cells [15–18]. However, a potential confounder in these studies is that the method used to detect isoforms has not been independently validated as accurate for scRNA-seq. We have demonstrated that Kallisto, Salmon, Sailfish and RSEM can accurately detect isoforms for scRNA-seq with high precision and recall, and we used Kallisto to reproduce the finding of previous studies that it is unusual for single cells to express multiple isoforms from a single cell [15–18]. In addition to being of biological relevance, this finding is important from a bioinformatics perspective, as it suggests that the problem of isoform quantification is made simpler by using scRNA-seq.

## Conclusions

For high quality simulated scRNA-seq datasets with a high number of reads/cell, RSEM, Kallisto, Salmon and Sailfish can accurately detect and quantify isoform expression. For simulated Drop-seq datasets with a low number of reads per cell, the precision of isoform detection is often less than 50%. As a consequence, isoform quantification cannot currently be sensibly performed on datasets in which only a few thousand reads have been sequenced per cell, even if a large number of cells have been sequenced. Isoforms with a high number of dropouts appear to be relatively challenging to quantify, possibly because such isoforms are often lowly expressed. We find that genes which express two isoforms in bulk RNA-seq predominantly express only one (or neither) isoform in equivalent scRNA-seq. In our benchmark of bulk RNA-seq, we discover the performance of most isoform quantification tools is slightly worse for scRNA-seq compared with bulk, but that the difference is small.

Taken together, our findings show that isoform quantification is possible with scRNA-seq for SMARTer and SMART-seq2 data. As single cells do not generally express all of the isoforms seen at the population level, scRNA-seq may eventually provide advantages over bulk RNA seq for isoform quantification by essentially deconvoluting the problem of isoform quantification. Future isoform quantification tools designed explicitly for scRNA-seq could improve on the performance of existing tools by being more conservative in calling isoforms as expressed, and by explicitly modelling the technical noise inherent to scRNA-seq.

## Methods

### Code Availability

The pipeline used to perform the RSEM simulation based BLUEPRINT and Kolodziejczyk et al. benchmarks can be found at https://github.com/jenni-westoby/Benchmarking_pipeline. The pipeline to perform the Splatter and Polyester simulation based benchmark can be found at https://github.com/jenni-westoby/Polyester_Simulations. The pipeline to perform the Drop-seq simulation based benchmark can be found at https://github.com/jenni-westoby/Drop-seq_pipeline. The pipeline to perform the bulk RNA-seq simulation based benchmark can be found at https://github.com/jenni-westoby/Benchmark_Bulk_Analysis. Instructions to perform isoform quantification on the real BLUEPRINT and Kolodziejczyk et al. bulk RNA-seq data can be found at https://github.com/jenni-westoby/Benchmark_paper_isoforms. The options and parameters passed to tools used to perform simulations and isoform quantification can be found at the above links.

A bug was encountered whilst using RSEM to simulate Drop-seq data. The bug was fixed and a pull request was made on the RSEM github page (https://github.com/deweylab/RSEM/pull/79).

The code and data used to generate the figures and supplementary figures used in this publication can be found at https://github.com/jenni-westoby/Benchmark_figures.

### Genomes

The Ensembl release 89 genome and transcriptome with 92 spike-in sequences developed by the External RNA Control Consortium (ERCC) appended were used wherever genome files in FASTQ format and/or transcriptomes in GTF format were required as input for tools in this study [47, 48]. The exception to this was when isoform quantification was carried out using the BLUEPRINT and Kolodziejczyk et al. bulk RNA-seq datasets. No ERCC spike-ins were added to these datasets, so the Ensembl release 89 genome and transcriptome without spike-ins appended were used to perform this analysis. To perform simulation and isoform quantification, RSEM produces a reference which includes a reference transcriptome in FASTQ format. This reference transcriptome produced by RSEM was used for isoform quantification tools which required a reference transcriptome in FASTQ format as input (See Github repository for code).

### Data Processing Prior to Analysis

Sequencing adaptors were trimmed from the Kolodziejczyk et al. and BLUEPRINT data using Cutadapt [49]. Reads from each cell in these datasets were aligned to the Ensembl genome release 89 using STAR [47, 50]. RSeQC was used to collect alignment quality statistics for each cell [51]. These statistics and the number of reads sequenced in each cell were used to remove low quality cells from each dataset (see Supplementary Figures S1 & S2). In addition, scater was used to plot the percentage of reads mapping to mitochondrial RNA and remove cells with greater than 10% of reads mapping to mitochondrial RNA [52] (See Supplementary Figures S1, S2, S3, S4, S10, S11, S13, S14).

Traditionally, Drop-seq data is not demultiplexed during gene level quantification [23]. However, with the exception of Kallisto, the tools used in this study cannot take multiplexed UMI data as input. When Kallisto does take multiplexed UMI data as input, it gives expression estimates for equivalence classes rather than for specific isoforms as output. Given that an anticipated issue with using Drop-seq for isoform quantification was that a UMI based method with 3’ coverage bias may not contain enough information to resolve between different isoforms from the same gene, it was decided that the performance of Kallisto when run in this mode would not be evaluated. Instead, the Shekhar et al. dataset was demultiplexed and RSEM was used to simulate a subset of the demultiplexed cells. The performance of eXpress, Kallisto, Sailfish, Salmon and RSEM was then evaluated in the simulated cells. The cell barcodes used to demultiplex the Drop-seq data were selected by following the instructions on the Drop-seq website to generate a gene expression matrix. The barcodes were extracted from the gene expression matrix and used to demultiplex the data. Further details can be found at https://github.com/jenni-westoby/Drop-seq_pipeline.

### Simulations

Two simulation methods were used in this study. The first method used to simulate single-cell RNA-seq data was RSEM. RSEM is an isoform quantification tool which makes use of a generative model and an expectation maximization algorithm to perform isoform quantification [4]. When performing isoform quantification, RSEM infers values for the latent variables in its generative model in addition to estimating isoform expression. To perform simulations, RSEM takes the inferred values of the latent variable and the expression estimates and uses them in its generative model to probabilistically simulate reads. As RSEM simulates reads, it counts where in the transcriptome each of the reads came from. RSEM thus simulates reads data for which it is known how highly expressed each isoform in the transcriptome is.

For each cell in the Kolodziejczyk et al. and the BLUEPRINT datasets that passed quality control and for each of the selected cells in the Drop-seq dataset, one RSEM simulation was performed. Isoform quantification was performed on each cell and the isoform expression estimates and inferred estimates for RSEM’s latent variables were used to perform the simulation. Consequently, each RSEM simulated cell used in this study was simulated using variables inferred from a real cell.

The second simulation method was based on two tools, Splatter and Polyester. Splatter is a simulation tool which takes an expression matrix of counts from a single-cell RNA-seq experiment as input and gives a simulated expression matrix of counts as output [53]. The Splatter package in fact contains six simulation methods. To select which performed best, data was simulated using the Lun, Lun2 and Simple simulation methods. The Splat simulation method was discounted as it was unable to simulate large enough expression matrices to account for the larger number of isoforms compared with genes, and the scDD method was discounted as it simulates differential expression, and no differential expression was expected. The BASiCS method had not been implemented at the time when the simulations were performed. Based on Splatter-generated graphs, the lun2sim method, inspired by a simulation method developed by Lun & Marioni [54] was selected as it bore the closest resemblance to the real data (see Figure S5).

The lun2sim method was used to simulate a matrix of counts based on an expression matrix of counts from the BLUEPRINT B lymphocytes generated by Kallisto [6]. The simulated expression matrix of counts was then given as input to Polyester, which simulated reads based on the lun2sim counts matrix. Simulations were performed both using Polyester’s uniform coverage model and using Polyester’s 3’ coverage bias model. The Splatter counts matrix was converted to a matrix of TPM values, which were used as the ‘ground truth’ for how highly expressed each isoform was in the Polyester simulated reads data.

### Post Simulation Data Processing

Reads from each cell in the datasets simulated by RSEM based on the Kolodziejczyk et al. and BLUEPRINT datasets were aligned to the Ensembl genome release 89 using STAR. RSeQC was used to collect alignment quality statistics for each cell. The alignment quality statistics and the number of reads for each simulated cell were used to remove low quality cells from each dataset (see Figures S3 & S4). Scater was used to plot the percentage of reads mapping to mitochondrial RNA and remove cells with greater than 10% of reads mapping to mitochondrial RNA.

### Bulk RNA-seq analysis

Prior to isoform quantification, RSeQC was used to remove rRNA mapping reads from the BLUEPRINT B lymphocyte bulk RNA-seq data. The code used to generate the isoform expression matrices used in the bulk RNA-seq benchmark can be found at https://github.com/jenni-westoby/Benchmark_Bulk_Analysis. The parameters used to run Kallisto on the real BLUEPRINT and Kolodziejczyk et al. datasets to determine how many isoforms were expressed per gene can be found at https://github.com/jenni-westoby/Benchmark_paper_isoforms.

### Statistics

The formula used to calculate the normalized root mean square error (NRMSE) is:

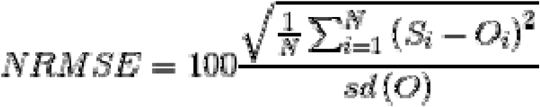

Where *N* is the number of isoforms that could have been simulated, S is the isoform expression estimates for the isoform quantification tool of interest, *O* is the ground truth expression estimates and *sd(O)* is the sample standard deviation of the ground truth expression estimates.

Prior to calculating the NRMSE, the ground truth and the isoform expression estimates were transformed using the formula:

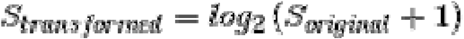

Where *S*_*original*_ was the original value of the ground truth or the expression estimate. This transformation reduces the impact of a small number of highly expressed isoforms on the value of the NRMSE.

## Declarations

### Availability of Data and Materials

The Kolodziejczyk et al. ES cell data was accessed from the ArrayExpress database (http://www.ebi.ac.uk/arrayexpress) using the accession number E-MTAB-2600, as described in the Kolodziejczyk et al. paper [40]. The Shekhar et al. Drop-seq data was accessed under GEO accession number <u>GSE81905</u>, as described in the Shekhar et al. paper [42]. The BLUEPRINT data was accessed under GEO accession number GSE94676.

### Author’s Contributions

JW, AFS and MH conceived the study and designed the experiments. JW carried out the experiments and wrote the manuscript. MS generated the BLUEPRINT B and T lymphocytes bulk and scRNA-seq data sets. MH and AFS supervised the experiments. All authors reviewed and approved the manuscript.

## Acknowledgements

We would like to thank Guillermo Parada, Vladimir Kiselev and Tallulah Andrews for their advice on developing pipelines. We would additionally like to thank Vladimir Kiselev for his advice on managing results data. We would like to thank Sarah Teichmann and Aleksandra A. Kolodziejczyk for scientific advice and experimental support on the preparation of cells for the C1 system. We would like to thank Russell Hamilton, Sudhakaran Prabakaran and Tallulah Andrews for their comments on the manuscript. AFS and MS were funded by BLUEPRINT (HEALTH-F5-2011-282510) and Wellcome Trust WT095606RR to AFS. MH was supported by the core funding provided to the Wellcome Trust Sanger Institute by the Wellcome Trust.

